# Environmental Consistency Modulation of Error Sensitivity During Motor Adaptation is Explicitly Controlled

**DOI:** 10.1101/528752

**Authors:** Guy Avraham, Matan Keizman, Lior Shmuelof

## Abstract

Motor adaptation, the adjustment of sensorimotor representations in face of changes in the environment, may operate at different rates. When human participants encounter repeated or consistent perturbations, their corrections for the experienced errors are larger compared to when the perturbations are new or inconsistent. Such modulations of error sensitivity were traditionally considered to be an implicit process that does not require attentional resources. In recent years, the implicit view of motor adaptation is challenged by evidence showing a contribution of explicit strategies to learning. These findings raise a fundamental question regarding the nature of the error sensitivity modulation processes. We tested the effect of explicit control on error sensitivity in a series of experiments, in which participants controlled a screen cursor to virtual targets. We manipulated environmental consistency by presenting rotations in random (low consistency) or random walk (high consistency) sequences, and illustrated that perturbation consistency affects the rate of adaptation, corroborating previous studies. When participants were instructed to ignore the cursor and move directly to the target, thus, eliminating the contribution of explicit strategies, consistency-driven error sensitivity modulation was abolished. In addition, delaying the visual feedback, a manipulation that affects implicit learning, did not influence error sensitivity under consistent perturbations. These results suggest that increases of learning rate in consistent environments are attributable to an explicit rather than implicit process in sensorimotor adaptation.

**Significant Statement**

When experiencing an error in a motor task (e.g., missing a basketball shot in a windy day), the motor system modifies its next action based on environmental consistency (how frequent the changes in wind’s direction and strength are). It is unknown whether this process is driven by an implicit and automatic process, or by an explicit process that employs cognitive strategies. We examined these possibilities in a simple visuomotor task by perturbing the feedback in each trial with different consistency levels, and manipulating the use of implicit and explicit processes. We found that participants increase their sensitivity to errors in consistent environments when employing explicit strategies, and do not change their behavior when the implicit process is operating alone.

## Introduction

A fundamental principle in motor learning is modularity. Even simple motor learning behaviors, like adaptation, are driven by multiple learning processes (Smith et al., 2006). A striking behavioral account for modularity in sensorimotor adaptation was demonstrated by the existence of implicit and explicit processes (Mazzoni and Krakauer, 2006; Taylor and Ivry, 2011; Haith and Krakauer, 2013; Taylor et al., 2014). The Implicit learning process refers to an automatic recalibration of the motor response to an experienced error, whereas the explicit learning process is the intentional update of aiming direction following a strategy to improve performance. These processes are also thought to be associated with different neural substrates; implicit learning depends on the cerebellum (Imamizu et al., 2000; Taylor et al., 2010; Galea et al., 2011; Schlerf et al., 2012; Kim et al., 2015; Morehead et al., 2017) whereas explicit learning is associated with cortical function (Taylor and Ivry, 2014; McDougle et al., 2016) and the dopaminergic system (Leow et al., 2012).

Despite the accumulation of results supporting this modularity, the functional roles of the implicit and explicit processes, and the interaction between them during adaptation, are still largely unknown. It was proposed that each learning process is driven by a different error signal; the implicit process is driven by sensory prediction errors, the difference between the expected and the actual sensory feedback, and the explicit process is driven by target error, the difference between the target and the feedback (Mazzoni and Krakauer, 2006; Taylor and Ivry, 2011; Shmuelof et al., 2012a; Reichenthal et al., 2016). This idea can explain the parallel operation of these processes during the time course of visuomotor adaptation (Taylor et al., 2014). However, it does not explain secondary influences on learning, such as modulations of *error sensitivity* – the change in the reaction to errors – that were reported for different error magnitudes (Criscimagna-Hemminger et al., 2010; Marko et al., 2012) and for different perturbation consistencies (Herzfeld et al., 2014).

Error sensitivity was shown to increase for small errors (Marko et al., 2012). However, this dependency is challenged by recent evidence showing invariance of the implicit process to error magnitude, i.e., that different errors lead to a constant and signed motor correction (Bond and Taylor, 2015; Morehead et al., 2017; Kim et al., 2018). The apparent error-sized dependent modulation of error sensitivity may therefore be an outcome of dividing the (fixed) correction by different error magnitudes.

Another contextual effect on error sensitivity is the consistency of the perturbation. Gonzalez Castro et al., 2014 and Herzfeld et al., 2014 have shown that humans adapt faster to perturbations that are consistent compared to perturbations that are random (Gonzalez Castro et al., 2014; Herzfeld et al., 2014). Importantly, sensitivity to error in these studies was measured for probe trials in which the experienced error was similar across the different consistency conditions, thereby controlling for the concern that modulation of error sensitivity was merely a normalization artifact that reflects differences in error magnitudes. The effect of consistency on adaptation pose an important question regarding its underlying mechanism; on the one hand, the increased learning rate for the consistent errors could be a result of an implicit error-sensitivity modulation in the cerebellum (Herzfeld et al., 2014; Hanajima et al., 2015), or alternatively, consistency can increase the awareness of the participant to the perturbation and thereby enhance the involvement of strategies.

In a series of visuomotor rotation experiments, we take a close look at the interaction between awareness and perturbation’s consistency. We report that the modulation of error sensitivity due to the consistency of the perturbation depends on explicit processes, and that manipulating implicit learning has no detectable effect on error sensitivity under consistent perturbations. Our results provide further support for the lack of contextual effects on implicit learning, and emphasize the crucial role of explicit control in sensorimotor learning.

## Methods

### Participants

Seventy-one healthy right-handed volunteers (aged 19-34 years; 44 females) participated in three experiments: 18 in Experiment 1, 31 in Experiment 2, and 22 in Experiment 3. All experiments were conducted after the participants signed an informed consent form approved by the Human Subjects Research Committee of Ben-Gurion University of the Negev, Be’er-Sheva, Israel.

### Experimental setup and task

Participants sat facing a computer monitor (resolution-1280 × 1024 pixels, dimensions: 37.7 × 30.1 cm), distant by ∼1.5 meters, and controlled a screen cursor by making pointing movements with the fist through flexion-extension and pronation-supination of the right wrist (Krakauer et al., 2006; Shmuelof et al., 2012b). Their right forearm rested within a stabilization device that prevented its supination. The cursor location on the screen was mapped to the position of a retroreflective marker attached to the knuckle of the index finger; it was calibrated such that a 1 cm deviation of the marker caused a 3.2 cm deviation of the cursor. The marker position was recorded at 100 Hz using three motion capture cameras (Qualisys AB, Sweden).

In all experiments, participants were requested to move the cursor to the target by performing a wrist pointing movement (Fig. 1). The start location, depicted as a circle in the center of the screen, and a gray target, 1.2 cm diameter and distant by 6.3 cm from the start location, were both presented on the screen throughout the trial. Each trial was initiated with the appearance of an orange cursor, 0.6 cm diameter, simultaneously with a presentation of a tone, signaling the participants to move to a start location, which was colored in blue. If participants remained in the start location for 1 sec, they received a ‘Go’ cue-both cursor and start location turned black, marking the onset of a fast out-and back movement with the goal of placing the reversal point on the target. To eliminate online feedback corrections, the cursor disappeared as soon as it traveled a radial distance of 10% of the distance to the target. 2 secs after the ‘Go’ cue, participants received performance feedback: the black cursor reappeared at the reversal point and the target changed colored either to green for target hits, or to red for misses. We considered a hit when the center of the cursor was distant by less than 0.65 cm from the center of the target (i.e., when the cursor’s center was inside the target). Participants could experience two types of trials: Contingent and Non-Contingent (Error Clamp). In Contingent trials, the location of the feedback cursor was contingent on participants’ movements, either veridical or rotated (to the clockwise or counterclockwise direction) with respect to movement direction. In Error Clamp trials, the cursor landed in a position that was non-contingent on participants’ movements; this position was rotated by 15^0^ clockwise or counterclockwise with respect to the target, and in both cases, the cursor appeared at the radius of the target. The purpose of the Error Clamp trials was to measure error sensitivity for a constant error size.

**Figure 1.**
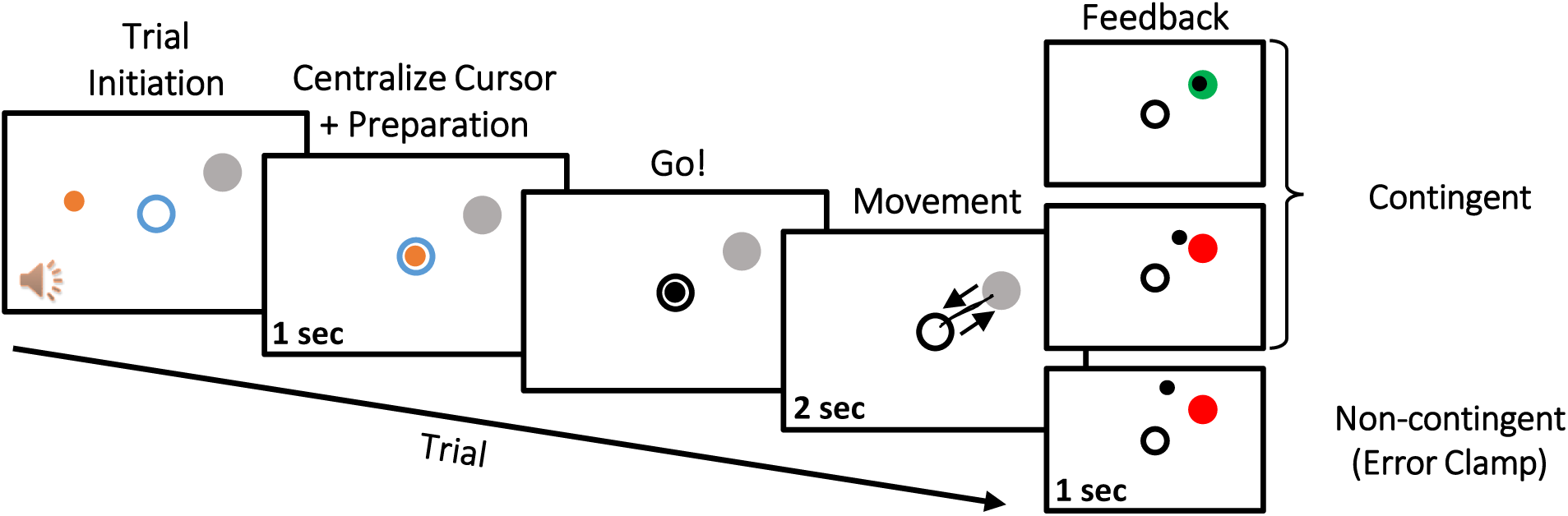
Experimental task. Trial initiation was marked by the appearance of a cursor (orange dot) and a presentation of an auditory tone. Participants were requested to move the cursor to a start location (blue circle). Following 1 sec in the start location, they received a ‘Go’ cue-both cursor and start location turned black, signaling participants to move to the target (filled gray circle). Participants were requested to perform a fast out-and-back movement, placing the reversal point on the target. The cursor was not presented during the movement. Movement directions (arrows) and an example of a movement path (black line) are schematically illustrated and were not presented to the participants. Trials ended with participants receiving performance feedback: a black dot reappeared at the reversal point and the target changed colored either to green for target hits, or to red for misses. In Contingent trials, the location of the feedback cursor was contingent on participants’ movements. In Non-contingent (Error Clamp) trials, the cursor landed at the radius of the target, in a predetermined position that was rotated by 15^0^ clockwise or counterclockwise with respect to the target.

### Experimental protocol

In all three experiments, participants did multiple experimental runs. Each run started with a baseline epoch with veridical visual feedback. This epoch was followed by an adaptation epoch in which the cursor was rotated with respect to the movement direction of the hand. Rotation magnitudes ranged from −30^0^ to 30^0^ in steps of 5^0^ (negative and positive values represent counterclockwise and clockwise, respectively). Runs were different by the schedule of the presented rotations, and each run comprised of one of two types of conditions that varied by consistency: Random and Random Walk (see Fig. 2 for illustration of each type). For the Random condition, the rotations were presented in a pseudorandom order, and changed between successive trials, such that the consistency, measured by lag-1 autocorrelation (Gonzalez Castro et al., 2014), is small (Table 1). For the Random Walk condition, the rotations varied from trial to trial according to a random walk algorithm: for each successive trial, the rotation changed by 5^0^, either clockwise or counterclockwise. The resulting perturbation function of the Random Walk condition had higher consistency than the Random perturbation function (Table 1).

**Table 1.** lag-1 autocorrelation (*R* (1)) values. A single value represents the *R* (1)of a perturbation schedule in a single run, i.e., the correlation coefficient between the rotation magnitudes in two successive trials. All values within each cell represent all possible runs in an experiment.

|  | Random | Random Walk |
| --- | --- | --- |
| **Experiment 1** | 0.25, 0.27, 0.18, 0.26 | 0.91, 0.91, 0.92, 0.91 |
| **Experiment 2** | 0.04 | 0.93 |
| **Experiment 3** |  | 0.91, 0.92, 0.89, 0.93 |

**Figure 2.**
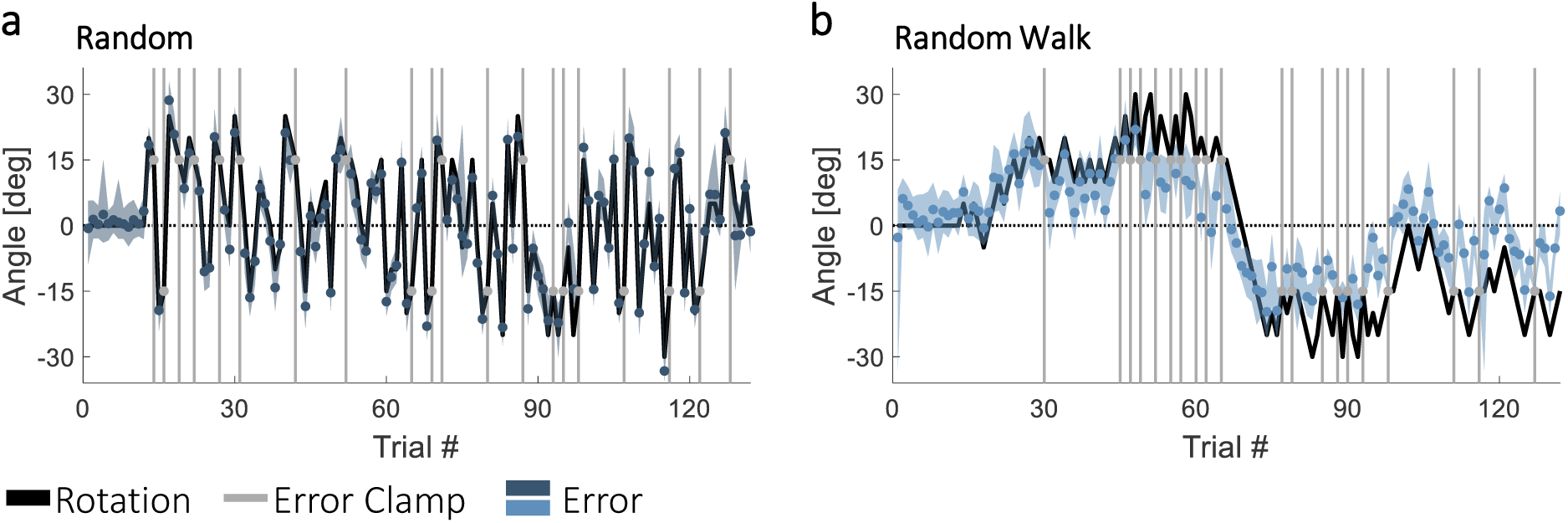
Experiment 1: Angular errors during Random and Random Walk sequences of visuomotor rotation. Time courses of the applied rotation (black lines) and errors (colored dots) during Random (***a***, dark blue) and Random Walk (***b***, light blue) perturbation sequences. Gray dots and vertical lines represent Error Clamp trials. Shading represents 95% confidence interval.

Most of the trials during the adaptation epoch were Contingent trials and some trials were Error Clamp trials (see below for exact percentages in each experiment). For the Random condition, these two types of trials were presented in pseudorandom and predetermined order. For the Random Walk conditions, the Error Clamp trials were only presented after contingent trials that constitutes of rotation size that was similar to the clamp size (15^0^), i.e., 10^0^-20^0^; the purpose of this design was to avoid disruption of the high consistency of the Random Walk perturbation schedule.

#### Experiment 1

Experiment 1 consisted of a single group of participants (N=18). All participants did four experimental runs. In each run, they moved to a different target that was presented in one of four locations: 45^0^, 135^0^, 225^0^ and 315^0^. Each run started with a baseline epoch of 12 trials, followed by the adaptation epoch that consisted of 120 trials. ∼17% of the trials in the adaptation epoch (20 trials) were Error Clamp trials. During the adaptation epoch of two of the four runs, for which targets were separated by 180^0^ from each other, participants experienced Random perturbation sequences, and during the other two runs, they experienced the Random Walk perturbation sequences (for each consistency condition, each participant experienced two of the four *R*(1)values in the associated cells in Table 1). The two conditions alternated in each experiment, and we counterbalanced the condition that was presented on the first run across participants. Across participants, all targets were associated with both conditions.

#### Experiment 2

Experiment 2 consisted of two groups of participants. All participants did two experimental runs, moving to a single target, presented at 45^0^. In each run, the baseline epoch consisted of 20 trials, and the adaptation epoch consisted of 220 trials. 20 trials in the adaptation epoch (∼9%) were Error Clamp trials. Unlike in Experiment 1, where Error Clamp trials could appear anytime during the adaptation epoch, here they were presented only after the second half of the run (after trial #120) to ensure that participants had sufficient exposure to the perturbations and their error sensitivity could be adjusted. During the adaptation epoch of one of the two runs, participants experienced the Random perturbation sequence, and during the other run, they experienced the Random Walk perturbation sequence. All participants experienced the same Random and Random Walk perturbation sequences (Table 1). The order of the conditions was counterbalanced across participants.

To examine the role of awareness in error sensitivity modulations, participants of one group (Ignore, N=15) were briefed about the perturbation and were requested to ignore the cursor and to move their hand directly to the target. Participants of the other group (Compensate, N=16) were not told about the rotation and were instructed to keep trying to hit the target with the cursor. Such instructions were previously shown to control for the contributions of implicit (Ignore group) and explicit (Compensate group) processes in sensorimotor learning (Welch, 1969; Morehead et al., 2017).

#### Experiment 3

Experiment 3 consisted of a single group of participants (N=22). All participants did four experimental runs. In each run, they moved to a different target that was presented in one of four locations: 45^0^, 135^0^, 225^0^ and 315^0^. The order of the targets was the same for all participants. In each run, the baseline epoch consisted of 12 trials, and the adaptation epoch consisted of the 120 trials. 20 trials in the adaptation epoch (∼17%) were Error Clamp trials. During the adaptation epoch of all four runs, participants experienced Random Walk perturbation sequences (Table 1), and were requested to always try to hit the target with the cursor.

Across runs, we manipulated the implicit process by imposing different delays between the moment of movement reversal and the feedback presentation. Within each run, the delay was either constant at 1,000 ms or 2,000 ms, or varied randomly between 600-1,500 ms or 1,600-2,500 ms in steps of 100 ms. All participants experienced all four types of delay schedules, but the order was randomized between participants. Across participants, all targets were associated with all delay schedules.

### Data analysis

The marker position (attached to the fist) was recorded throughout the experiment and sampled at 60 Hz. It was analyzed offline using a custom-written MATLAB code (The MathWorks, Natick, MA, RRID: <u>SCR_001622</u>). To measure the movement reversal of the marker, for each trial, we identified the first sample (i) in which the movement amplitude was smaller than the previous sample (*i* − 1). The movement reversal was defined as the location of the marker on sample *i* − 1. We defined the hand’s movement angle (*MA*) as the angle between the imaginary lines connecting the movement origin to the movement reversal and to the target. We calculated the directional error (*e*_*n*_) at trial *n* as the angular difference between the feedback and the target. For each error, we measured learning from error (*LE*), as the change in *MA* from the trial in which the error was experienced to the next trials:

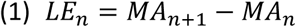

Error sensitivity (*ES*) was calculated as:

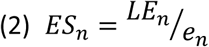

We calculated learning from error for all trials. Error sensitivity was obtained separately for all 15^0^ Error Clamp trials and for all trials in which the experienced absolute error was between 10^0^-20^0^ (Error Clamp and Contingent trials).

Learning from error functions were obtained by sorting errors experienced in each condition for each participant based on magnitude, binning them to 12 bins, and averaging the experienced errors and the change in hand angle within each bin. Then, we calculated the between-participant mean and standard error of the experienced errors and the change in hand angle for each bin.

We quantified the learning from error by fitting a linear function to the change in hand angle with respect to the error size for each participant and condition. We used the slope of the fitted linear function as a measure for learning from error across all of the experienced error magnitudes. Greater absolute slope values indicate higher error sensitivity.

### Statistical analysis

Statistical analyses were performed using custom-written MATLAB functions, the MATLAB Statistics Toolbox, and IBM SPSS (RRID: <u>SCR_002865</u>). We used the Lilliefors test to determine whether our measurements were distributed normally (Lilliefors, 1967). In the repeated-measures ANOVA models, we used Mauchly’s test to examine whether the assumption of sphericity was met. For the factors that were statistically significant, we performed planned comparisons and corrected for familywise error using the Bonfferoni correction. We denote the Bonfferoni-corrected *p* values as *p*_*B*_.

The statistical analyses for all three experiments were done on the following measures: (1) mean error sensitivity across all 15^0^ Error Clamp trials, (2) all 10^0^-20^0^ error trials, and (3) the slope of the learning from error function, calculated separately for each participant and each condition.

To examine the influence of consistency on each of the above error sensitivity measures in Experiment 1, for each participant, we pulled together the data from two runs of the same consistency condition (Random and Random Walk), and calculated each participant’s error sensitivity measures for each consistency condition. We used a two-tail paired-sample *t* test to examine whether the difference in each measure between the Random and the Random Walk conditions are statistically significant.

To examine the effects of explicit strategies on error sensitivity modulation in environment with different levels of consistency in Experiment 2, we calculated error sensitivity measures for each consistency condition. For each measure, we fitted a two-way mixed-effect ANOVA model, with the measure as the dependent variable, one between-participants independent factor (Strategy: two levels, Ignore and Compensate), and one within-participant independent factor (Consistency: two levels, Random and Random Walk).

To examine the effects of delayed feedback (modulation of implicit adaptation) on error sensitivity, we calculated each participant’s error sensitivity measures for each delay condition. For each measure, we fitted a four-way repeated-measures ANOVA model, with the measure as the dependent variable, and one within-participant independent factor (Delay: four levels, 600-1,500, 1,000, 1,600-2,500, and 2,000 ms).

Throughout this paper, statistical significance was set at the *p* < 0.05 threshold.

## Results

### Experiment 1: Consistency of the perturbation increases error sensitivity

A group of participants (N=18) experienced both Random and Random Walk schedules of visuomotor rotations on different experimental runs (Fig. 2). During the Random condition, the rotations were presented in a pseudorandom order, changing between successive trials, such that the consistency, measured by lag-1 autocorrelation (Gonzalez Castro et al., 2014), is small (*mean R*(1)= ∼0.24, see methods for further details). During the Random Walk condition, the rotations varied from trial to trial according to a random walk algorithm, resulting a perturbation function with higher consistency (*mean R*(1)= ∼0.91) than the Random perturbation function.

The time courses of the mean directional error (Fig. 2) suggests that participants adapted to some degree to the Random Walk perturbation; this is evident by the gradual decrease with respect to the rotation size, especially during the late stage of the adaptation epoch (Fig. 2b). The Random perturbation masks any improvement in performance (Fig. 2a).

During the Random Walk condition, participants showed higher sensitivity to errors than during the Random condition. To measure error sensitivity, we normalized the change in hand angle by the error size. For Error Clamp trials (trials in which the feedback cursor was presented 15^0^ away from the target, irrespective of the performance of the participant), despite the apparent increase in error sensitivity for the random walk group, this effect did not reach significance [*t* (17) = 1.40, *p* = 0.179, Fig. 3a]. However, computing error sensitivity for all the trials in which the experienced error size was within a range of 10^0^-20^0^, which allowed a more robust estimation of error sensitivity within each participant, reveals a higher sensitivity to these errors in the Random Walk condition than in the Random condition [*t* (17) = 2.66, *p* = 0.016, Fig. 3b]. We also examined the trial-by-trial change in hand angle as a function of the error size across the entire range of the experienced errors (learning from error, Fig. 3c). This analysis reveals that the absolute slope of the learning from error function for the Random Walk schedule is higher than for the Random perturbation [*t* (17) = 2.88, *p* = 0.010], suggesting that participants apply bigger corrections for the experienced error during the former than the latter condition. Additionally, participants experienced smaller errors during Random Walk than during Random conditions. The projections of the curves on the abscissa indicate that the errors experienced during the Random Walk condition has a narrower distribution than the errors experienced during the Random condition (as a result of the increased error sensitivity and the consistent perturbation schedule that allowed for a gradual reduction in error size). These results are in agreement with the previous reports that sensitivity to errors in sensorimotor tasks is higher as the consistency in the environment increases (Gonzalez Castro et al., 2014; Herzfeld et al., 2014).

**Figure 3.**
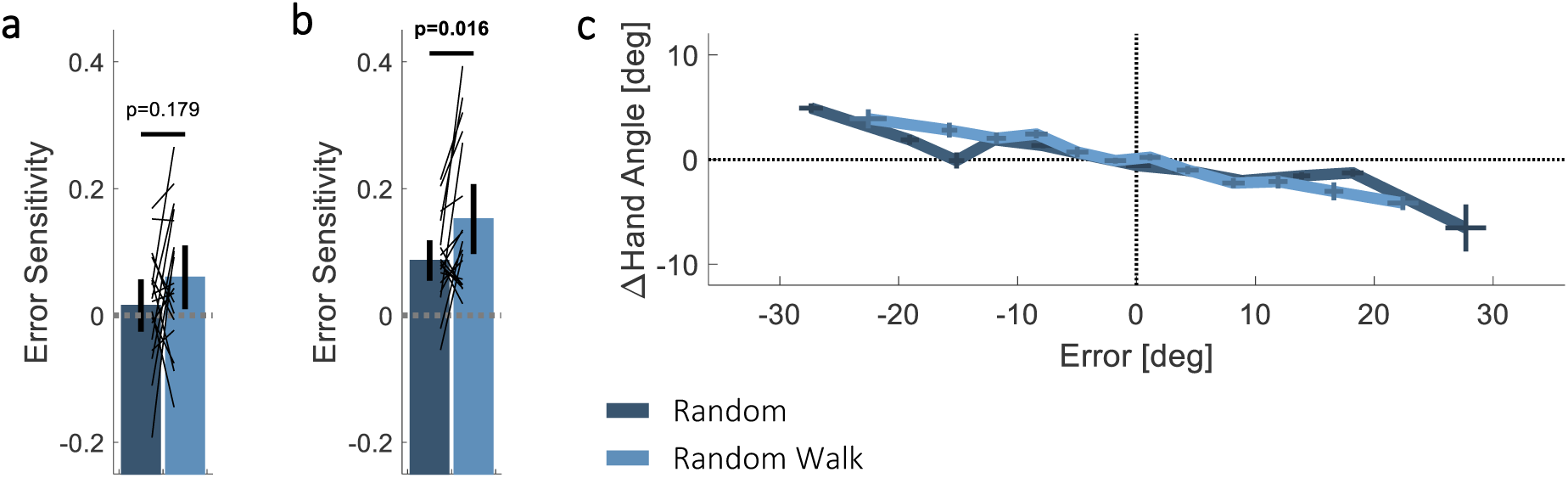
Experiment 1: Error sensitivity is higher for Random Walk than for Random sequences of visuomotor rotation. (***a, b***) Between-participants mean error sensitivity for each of the Random (dark blue) and Random Walk (light blue) conditions, computed for Error Clamp trials (15^0^ errors, ***a***) and for 10^0^-20^0^ error trials (***b***). Error bars represent 95% confidence interval. Thin black lines represent data of individual participants. Significant p-value is bolded. (***c***) Mean change in hand angle as a function of the experienced error for the Random and Random Walk perturbation sequences. The trials were binned by error size for each participant. The centers of the crosses represent the mean hand angle for the mean error of each bin, and the vertical and horizontal lines of the crosses represent between-participant standard error.

### Experiment 2: Increased error sensitivity in consistent environments depends on explicit strategies

The higher sensitivity to errors for the Random Walk condition with respect to the Random condition in Experiment 1 could have been driven by implicit changes in responses to the errors, by a modulation of explicit control strategies, or both. We examined these possibilities in Experiment 2.

Two groups of participants performed the same visuomotor task as in Experiment 1. Both groups experienced both Random (*R* (1)= 0.04) and Random Walk (*R* (1)= 0.93) perturbation schedules in different experimental runs. To examine the role of awareness in error sensitivity modulations, participants in the Ignore group (N=15) were requested to ignore the feedback and to move their hand directly to the target, whereas participants in the Compensate group (N=16) were instructed to keep trying to reach the target with the cursor.

Error sensitivity was the highest when participants were explicitly compensating for the Random Walk perturbation schedule. Statistical analyses of error sensitivity in the Error Clamp trials (15^0^ errors, Fig. 4a) and from the 10^0^-20^0^ error trials (Fig. 4b) revealed a significantly higher error sensitivity in the Compensate than in the Ignore group (Strategy main effect: 15^0^ errors-*F* (1,29) = 7.01, *p* = 0.013; 10^0^-20^0^ errors-*F* (1,29) = 19.15, *p* = 1.43 × 10^−4^), suggesting that the Ignore manipulation suppressed error corrections. Similar to Experiment 1, the participants increased their sensitivity to errors when faced with Random Walk rather than Random perturbation schedule in the 10^0^-20^0^ error trials (Consistency main effect: 15^0^ errors-*F* (1,29) =2.02, *p* = 0.166; 10^0^-20^0^ errors-*F* (1,29)=12.20,*p* =0.002). Most importantly, we found an interaction effect between strategy and consistency influences on error sensitivity modulation (Strategy-Consistency interaction effect: 15^0^ errors-*F* (1,29) = 5.58, *p* = 0.025; 10^0^-20^0^ errors-*F* (1,29) = 5.95, *p* = 0.021). While the Ignore group did not exhibit any change in error sensitivity between the Random and Random Walk conditions (15^0^ errors-*p*_B_ = 0.518; 10^0^-20^0^ errors-*p*_B_ = 0.469), the Compensate group had a significantly higher error sensitivity during the Random Walk condition than during Random condition (15^0^ errors-*p*_B_ = 0.011; 10^0^-20^0^ errors-*p*_B_ = 1.95 × 10^−4^). In addition, a comparison between the groups during the Random Walk condition revealed a significantly higher error sensitivity in the Compensate group with respect to the Ignore group (15^0^ errors-*p*_B_ = 0.007; 10^0^-20^0^ errors-*p*_B_ = 2.66 × 10^−4^).

**Figure 4.**
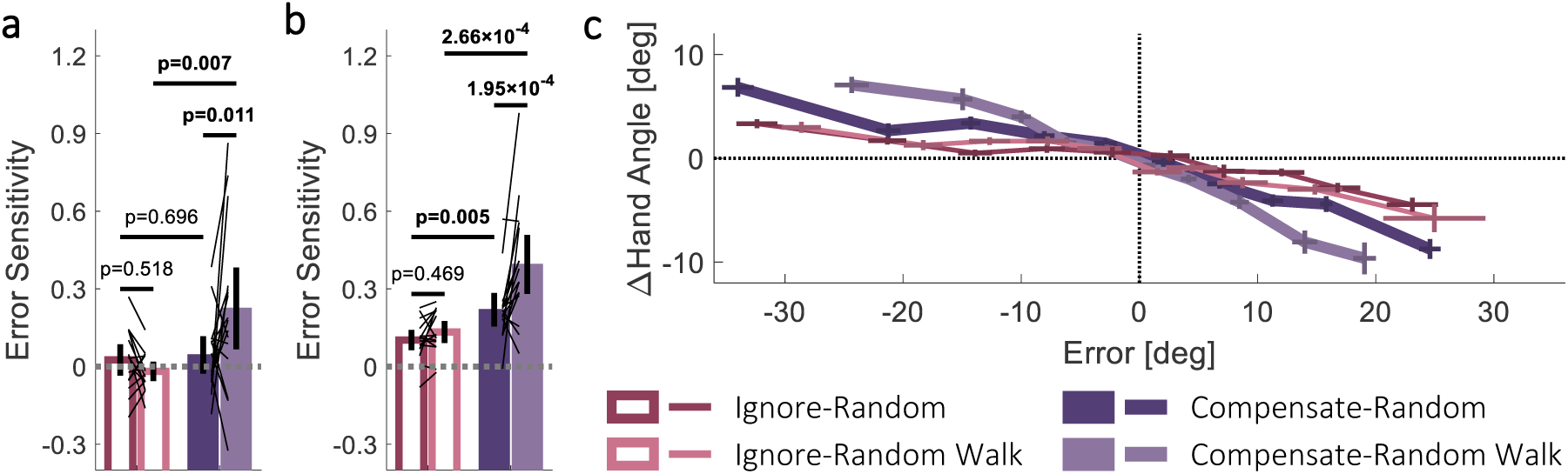
Experiment 2: Error sensitivity is higher for Random Walk than for Random sequences of visuomotor rotation when participants compensate rather than ignore the perturbations. (***a, b***) Between-participants mean error sensitivity for each of the Ignore (hollowed pink bars) and Compensate (filled purple bars) groups in each of the Random (dark color) and Random Walk (light color) perturbation sequence, computed for Error Clamp trials (15^0^ errors, ***a***) and for 10^0^-20^0^ error trials (***b***). Error bars represent 95% confidence interval. Thin black lines represent data of individual participants. Significant p-values are bolded. (***c***) Mean change in hand angle as a function of the experienced error for the Ignore (thin lines) and Compensate (thick lines) groups in each type of perturbation sequence. The trials were binned by error size for each participant. The centers of the crosses represent the mean hand angle for the mean error of each bin, and the vertical and horizontal lines of the crosses represent between-participant standard error.

The essential role of explicit strategies on error sensitivity modulation in consistent environments is also demonstrated by the analysis of the trial-by-trial change in hand angle across all experienced error magnitudes (Fig. 4c). The absolute slopes of the learning from error functions were significantly higher in the Compensate than in the Ignore group (Strategy main effect: *F* (1,29) = 23.75, *p* = 3.60 × 10^−5^), and during the Random Walk than the Random condition (Consistency main effect: *F* (1,29) = 20.29, *p* = 1.00 × 10^−4^). The difference between the groups increased when they experienced Random Walk compared to Random perturbation schedules (Strategy-Consistency interaction effect: *F* (1,29) = 9.76, *p* = 0.004). While the absolute slopes of learning from error functions of the Ignore group are comparable between the Random and Random Walk conditions (*p*_B_ = 0.344), the learning from error function of the Compensate-Random Walk condition has a larger absolute slope than the slope of the Compensate-Random function (*p*_B_ = 6.63 × 10^−6^). Furthermore, the abscissa projection of the learning from error function of the Compensate-Random Walk condition is narrower than all other functions indicating that during this condition, participants experienced the smallest distribution of errors. Overall, these results suggest that the increase in error sensitivity in consistent environments depends on the use of an explicit strategy.

### Experiment 3: modulations of feedback delay does not influence error sensitivity

The observed enhancement of error sensitivity in the Random Walk condition in the Compensate group in Experiment 2 is in line with the idea that the use of explicit strategies is required for modulating error sensitivity in consistent environments. In addition, the absence of difference in error sensitivity between the consistency conditions in the Ignore group suggests that implicit processes do not contribute to changes in error sensitivity. We verified the latter conclusion in Experiment 3.

One group of participants (N=22) experienced Random Walk perturbation schedules across different experimental runs (*R*(1)= ∼0.91). The participants were requested to always try to hit the target with the cursor. Across runs, we manipulated the implicit process by imposing different delays between the movement and the feedback. Delayed feedback was previously shown to attenuate adaptation (Kitazawa et al., 1995) through implicit processes (Brudner et al., 2016; Parvin et al., 2018). The magnitudes of the delays ranged between 600-2,500 ms. Within each run, the delay was either constant (1,000 or 2,000 ms), or variable (600-1,500 ms or 1,600-2,500 ms).

The delay of the feedback did not affect error sensitivity. We did not find statistically significant differences in error sensitivity for either Error Clamp (15^0^ errors, Fig. 5a, Delay main effect: *F* (3,63) = 0.78, *p* = 0.510) or 10^0^-20^0^ error trials (Fig. 5b, Delay main effect: *F* (3,63) = 0.29, *p* = 0.835). In addition, during all four delay conditions, participants exhibited similar learning from error functions that had comparable slopes (Fig. 5c, Delay main effect: *F* (3,63) = 0.82, *p* = 0.490). These results suggest that under random walk perturbation, suppression of the implicit process by increasing feedback delay does not affect error sensitivity, and therefore provide further support for the sole contribution of explicit control to enhanced error sensitivity in consistent environments.

**Figure 5.**
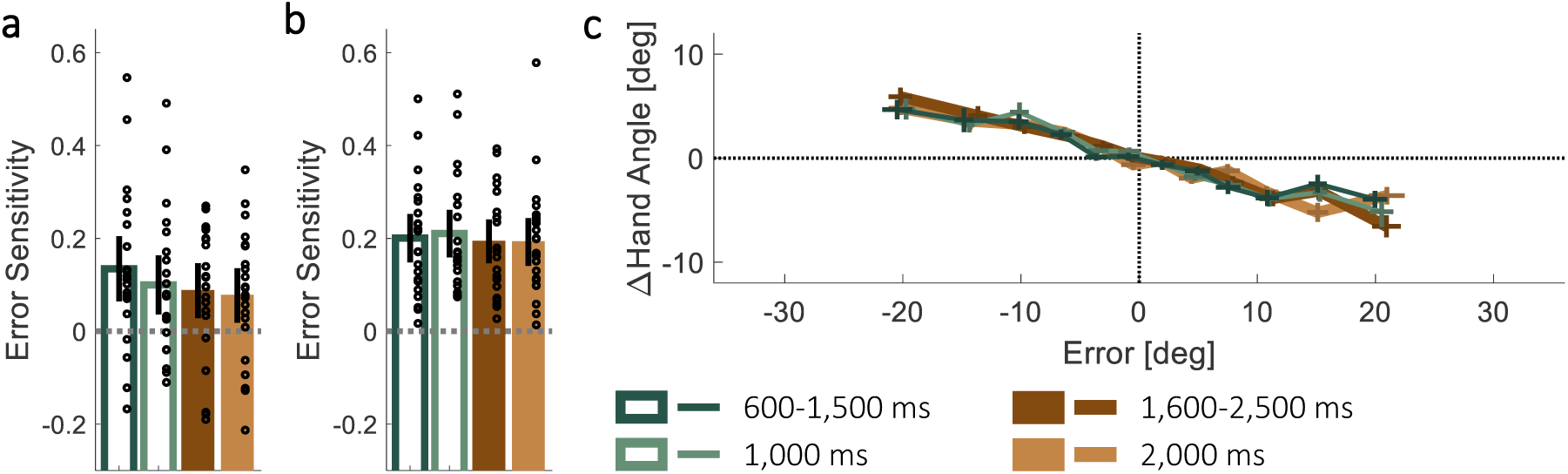
Experiment 3: Error sensitivity for Random Walk is comparable between different magnitudes of feedback delays. (***a, b***) Between-participants mean error sensitivity for feedback delays of 600-1,500 ms (hollowed dark green bar), 1,000 ms (hollowed light green bar), 1,600-2,500 ms (filled dark brown bar) and 2,000 ms (filled light brown bar), computed for Error Clamp trials (15^0^ errors, ***a***) and for 10^0^-20^0^ error trials (***b***). Error bars represent 95% confidence interval. Black circles represent data of individual participants. (***c***) Mean change in hand angle as a function of the experienced error for each of the short (thin lines) and long (thick lines) delay conditions. The trials were binned by error size for each participant. The centers of the crosses represent the mean hand angle for the mean error of each bin, and the vertical and horizontal lines of the crosses represent between-participant standard error.

## Discussion

Environmental consistency is considered to be an important factor in sensorimotor learning (Burge et al., 2008). We corroborated this here by demonstrating that sensitivity to errors is higher when experiencing random walk sequences of visuomotor perturbations compared to random sequences. Nevertheless, we show that consistency by itself is not a sufficient condition for increasing learning rates. When instructed to ignore the perturbations, a manipulation that suppresses explicit processes and thereby reveal the contribution of implicit learning processes (Mazzoni and Krakauer, 2006; Morehead et al., 2017), our participants did not exhibit an increase in error sensitivity in a consistent environment. Furthermore, variation of the delay of the feedback, a manipulation that typically influences implicit adaptation, yielded comparable levels of sensitivity to errors under random walk conditions, supporting the contribution of explicit learning processes to adaptation in consistent environments. Overall, our results suggest that the sensorimotor system increases error sensitivity in consistent environments due to the contribution of explicit strategies rather than by influencing the rate by which internal models are implicitly updated.

Previous examinations of consistency effects on sensorimotor learning largely ignored the roles of conscious awareness of the perturbations. Nevertheless, recent evidence illustrated that the sensitivity function of the explicit learning process to different magnitudes of visuomotor rotations is influenced by environmental consistency (Hutter and Taylor, 2018). Interestingly, the observed effect was non-monotonic: sensitivity increased from inconsistent to low-consistent perturbations, but decreased as consistency increased farther. Importantly, the sensitivity in Hutter and Taylor, 2018, was examined as a function of the perturbation size rather than the experienced visual error, and the distribution of the latter varies at different levels of consistency (Hutter and Taylor, 2018), which may explain the non-monotonicity behavior of the sensitivity function.

The main studies that examined consistency effects on error sensitivity used force field perturbations (Gonzalez Castro et al., 2014; Herzfeld et al., 2014). While it is difficult to isolate the contribution of explicit strategies to the compensation for force feedback, previous results with this paradigm may address another central question in motor learning: Do the environmental manipulations affect the sensitivity to error directly, or elicit other explicit strategies, such as a recall of a previous successful action? Gonzalez Castro et al., 2014 showed that if participants are suddenly exposed to a perturbation that is opposite to the trained perturbations, they react to it in the same way that they reacted to the trained perturbations, i.e., they apply a negative and inappropriate correction (Gonzalez Castro et al., 2014). This behavior reflects an involvement of a process other than error sensitivity modulation, such as a recall of previous correct responses (Haith and Krakauer, 2014). Our results support that notion by showing that consistency effects are mediated by explicit control.

Which strategy underlies the faster adaptation to consistent perturbations? A recent study provided evidence for distinct cognitive strategies during sensorimotor learning of visuomotor rotation: A recall of stimulus-response contingencies and parametric computation of error correction (McDougle and Taylor, 2019). The former relates to a fast process that utilizes memories of acquired associations between stimuli and responses, whereas the latter is a process that computes the aiming direction by means of mental rotation based on the experienced target error (Taylor and Ivry, 2011). We speculate that in the context of a constantly changing perturbations, the consistency in the random walk condition may enhance the mental rotation strategy of the participants. Possibly, a prolonged practice with a consistent perturbation would enable a caching process of stimulus response associations (Huberdeau et al., 2017) mediated by an increase in movement repetitions (Huang et al., 2011; Mawase et al., 2018).

The link between error sensitivity and explicit sensorimotor learning is also studied in the context of savings, i.e., the faster learning upon re-exposure to the same perturbation. Herzfeld et al., 2014 explained savings as a change in error sensitivity in face of an error that was previously encountered (Herzfeld et al., 2014). More recently, savings was explained as a retrieval of an explicit aiming strategy that was previously associated with a better performance (Haith et al., 2015; Morehead et al., 2015). Interestingly, despite previous evidence for the necessary role of both error-driven adaptation and movement repetitions in inducing savings (Huang et al., 2011), Leow at al., 2016 challenged this view, suggesting that the experience of previously encountered errors is sufficient as long as the perturbation is adequately stable (Leow et al., 2016). Our results are in line with this idea and suggest that savings is a product of enhanced sensitivity to an error that was processes explicitly due to the experience of consistent perturbations. Thus, savings can be viewed as an exemplar for more general phenomena of error correction facilitation that is driven by strategic processes. This conjecture also leads to the prediction that the neural substrates of savings that are localized to the motor cortex (Li et al., 2001; Landi et al., 2011) and basal ganglia circuits (Leow et al., 2012; Ruitenberg et al., 2018) will also be involved in the increased corrections for consistent perturbations. Nonetheless, savings and reaction to consistency may differ in terms of the processes that underlie their formation; while savings possibly depends on encoding of successful reactions to distinct errors (Huang et al., 2011; Huberdeau et al., 2015), consistency detection requires a continuous estimation of the ability to reduce external errors (Baddeley et al., 2003; Gonzalez Castro et al., 2014).

The current study supplements the growing body of literature that emphasizes the important contribution of explicit strategies to sensorimotor learning. Whereas the implicit process is highly rigid, the explicit process can be flexibly tuned to task demands, e.g., by scaling the correction to the error size (Bond and Taylor, 2015), and as mentioned above, by adjusting the reactions according to previous experience (Haith et al., 2015; Morehead et al., 2015; Huberdeau et al., 2017). Furthermore, age-related declines in motor learning were recently shown to be associated with both behavioral and neural changes in the explicit memory system (Vandevoorde and Xivry, 2018; Wolpe et al., 2018). Here we show that interaction between environmental consistency and motor learning is related to the modulations of explicit strategies rather than to the modulation of implicit adaptation.

Error clamp trials were previously used to probe changes of sensorimotor representations in face of dynamic (Scheidt et al., 2000; Avraham et al., 2017) and kinematic (Shmuelof et al., 2012a; Morehead et al., 2017) perturbations. In our design, each experimental run consisted of multiple error clamp trials with 15^0^ error size, aiming to provide sufficient recurrences for calculating error sensitivity for a constant error. Surprisingly, in some of our experiments, analysis of the error clamp trials yielded no change in error sensitivity (Figs. 3a and 4a). As these trials include feedback that is non-contingent on participants’ movements, the fact that they were presented multiple times within a continuously changing environment may have devaluated the relevancy of the associated error size, which in turn was discounted by the motor system (Wei and Körding, 2009). Therefore, we also examined error sensitivity for trials in which the experienced error was similar in size to the error experienced during error clamp trials (10^0^-20^0^). These analyses revealed that error sensitivity was indeed modulated in all our conditions (e.g., Figs. 3b and 4b). Importantly, the analysis results of the error clamp trials show that error sensitivity is largest under consistent environments and an explicit instruction to compensate for the errors, and thus, they are in line with the main conclusion of this study.

The association between consistency and explicit strategies is mediated by the detection of the consistency. Consistency estimation requires monitoring both the perceived errors and the reaction to these errors, but could also be approximated by memorizing the history of error magnitudes alone (by either averaging recent errors or monitoring their trial-by-trial changes). The use of explicit strategies may contribute to the consistency estimation process by increasing the awareness to the error (Johnson et al.,2002), enhancing the notion of agency (Parvin et al., 2018), or facilitating the generation of memories for the experienced errors (Morehead et al., 2015). Understanding the benefits and the driving signals of explicit learning strategies is essential for improving motor learning in general, and specifically, could improve the outcomes of rehabilitative treatments.

